# Flat clathrin lattices are linked to metastatic potential in colorectal cancer

**DOI:** 10.1101/2023.01.12.520576

**Authors:** Charlotte Cresens, Guillermo Solis-Fernandez, Astha Tiwari, Rik Nuyts, Johan Hofkens, Rodrigo Barderas, Susana Rocha

## Abstract

Clathrin assembles at the cells’ plasma membrane in a multitude of clathrin-coated structures (CCSs). Among these are flat clathrin lattices (FCLs), alternative clathrin structures that have been found in specific cell types, including cancer cells. Here we show that these structures are also present in different colorectal cancer (CRC) cell lines, and that they are extremely stable with lifetimes longer than 8 hours. By combining cell models representative of CRC metastasis with advanced fluorescence imaging and analysis, we discovered that the metastatic potential of CRC is associated with an aberrant membranous clathrin distribution, resulting in a higher prevalence of FCLs in cells with a higher metastatic potential. These findings suggest that clathrin organization might play an important yet unexplored role in cancer metastasis.

## Introduction

Clathrin-mediated endocytosis (CME) is essential for the selective internalization of cell surface receptors in mammalian cells. The main actor in this process is clathrin, a triskelion-shaped protein consisting of three heavy and three light chains, which binds cargo molecules at the plasma membrane. Through the recruitment of additional triskelions, a honeycomb-like lattice consisting of hexagons and pentagons assembles. According to the CME model for *de novo* vesicle formation, this lattice can form a shallow invagination called clathrin-coated pit (CCP). Assisted by adaptor proteins, a CCP can bend further, increasing its curvature and the depth of the invagination, until a spherical clathrin-coated vesicle (CCV) is released into the intracellular environment.

In agreement with the general CME outline, a variety of clathrin-coated structures (CCSs) have been observed, ranging from nearly planar lattices or slightly curved CCPs to domed structures or highly curved CCVs (1). However, alternative clathrin structures that do not fit this model have also been described, including flat clathrin lattices (FCLs, also termed clathrin plaques) (2–6). These structures are reported to be planar, heterogeneous and irregularly shaped non-classical CCSs that are too large to be resolved into a single diffraction-limited CCV (2–6). FCLs are functional endocytic structures that can release CCVs (4,7–9), but are more stable: while classical CCSs remain at the plasma membrane for about 30-90 seconds (10–12), FCLs are significantly longer-lived, with reported lifetimes ranging from 2-10 minutes to more than 1 hour (5,6,13). FCLs have exclusively been found in specific cell types, including several cancer cell lines (6,14), but a potential link between the presence of non-classical CCSs and cancer progression has not yet been investigated.

Previous research has shown that endocytosis is dysregulated in cancer (15–18) and that several proteins involved in CME are implicated in tumor cell migration, invasion and metastasis (19). By using subcellular proteomic analysis, we have previously established that proteins involved in vesicle trafficking, including in CME, are among the most dysregulated complexes in highly metastatic colorectal cancer (CRC) cells compared to poorly metastatic CRC cells (20,21).

In this work, we combined two isogenic cell models representative of CRC metastasis (KM12 and SW) with advanced fluorescence imaging to investigate whether cellular clathrin distribution is related to metastatic potential. The KM12 model consists of the poorly metastatic KM12C cell line obtained from a patient’s primary CRC tumor, and the liver metastatic KM12SM and liver and lung metastatic KM12L4a cell lines, obtained through successive passages in nude mice (22,23). The second model (namely SW) consists of the SW480 and SW620 pair of cell lines. The poorly metastatic SW480 cell line was obtained from a patient’s primary CRC tumor and the metastatic SW620 cell line was derived from lymph node metastasis in the same patient (24,25). Previous research reported a good correlation between findings in these cell models and patient samples (21,26–32), indicating that they quite adequately mimic the different subtypes of CRC patients and the critical steps of CRC metastasis. We investigated clathrin organization and dynamics using standard confocal microscopy, and we quantified detailed clathrin topology using super-resolution PALM-TIRF microscopy and customized analysis algorithms. Our results show that extremely stable alternative CCSs, identified as FCLs, are far more prevalent in highly metastatic cells compared to their poorly metastatic counterparts. This suggests that aberrant clathrin organization could potentially be a key player in the process of cancer metastasis.

## Results

### 1. Stable FCLs are present in metastatic CRC cells

To investigate whether alternative CCSs are present in the KM12 cell model, we imaged the clathrin distribution at the ventral membrane of cells transiently transfected with clathrin light chain A fused to EYFP (EYFP-CLTA). Confocal images indicate that poorly and highly metastatic cells display remarkable differences in clathrin topology (Fig 1). In the poorly metastatic KM12C cell line, clathrin predominantly appears as diffraction-limited dot-like structures, representing classical CCSs that fit the general CME model. In comparison, the two metastatic KM12SM and KM12L4a cell lines present an aberrant clathrin organization with both classical CCSs as well as larger, more elongated CCSs with longitudinal cross-sections reaching a few micrometers, which resemble FCLs. Previous reports have shown that transient overexpression of the CLTA plasmid does not influence clathrin topology (33), and our results support this finding since immunolabeling of endogenous clathrin heavy chain 1 (CLTC) resulted in a comparable topology (Fig S1).

**Figure 1:**
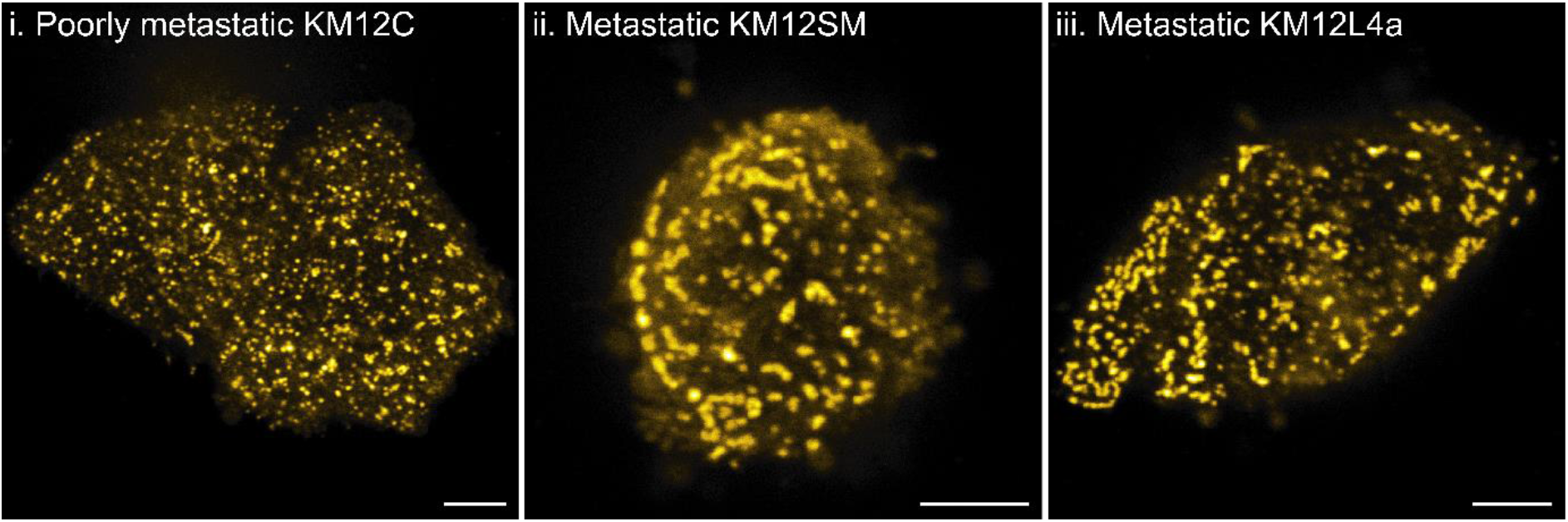
Clathrin topology imaged with confocal fluorescence microscopy. Representative images of transiently transfected EYFP-CLTA visualized at the ventral plasma membrane (bottom, in contact with the glass coverslip) of poorly metastatic KM12C (i), metastatic KM12SM (ii) and metastatic KM12L4a (iii) CRC cells. Scale bars 5 μm.

To confirm that the observed elongated CCSs are indeed FCLs, we monitored clathrin dynamics in metastatic KM12SM cells for longer time intervals. As shown in Figure 2A, classical dot-like CCSs are typically formed and internalized within 1-2 minutes, in line with their dynamic behavior during CME. In contrast, nearly all alternative CCSs remained present for longer periods of time and could still be observed after 8 hours in both the metastatic KM12SM and KM12L4a cell lines (Fig 2B and Fig S2). This extended lifetime supports the conclusion that these structures are FCLs. However, to the best of our knowledge, imaging of clathrin in FCLs for such prolonged periods of time has not yet been reported.

**Figure 2:**
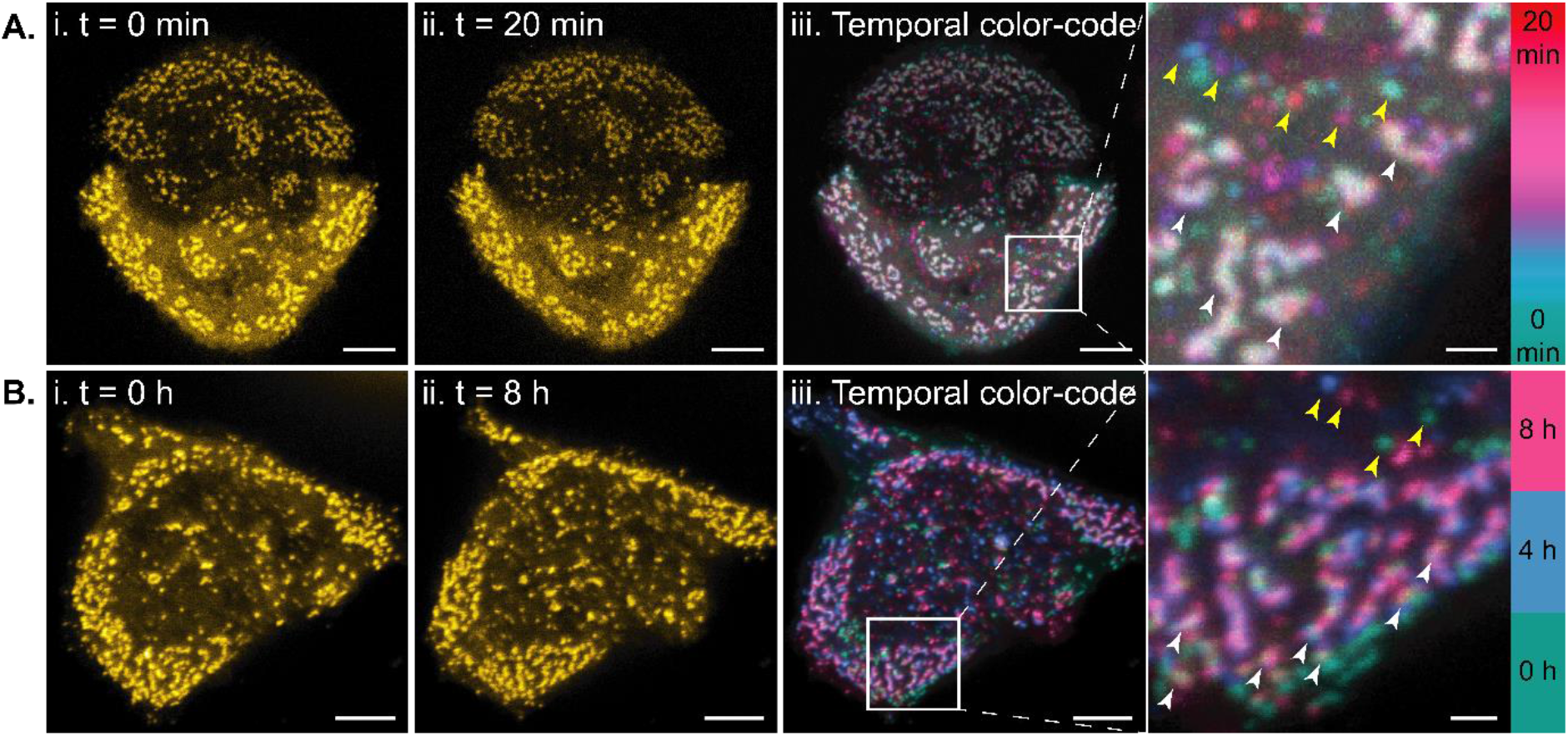
Clathrin dynamics at the ventral membrane of metastatic KM12SM cells transiently transfected with EYFP-CLTA, imaged by time-lapse confocal microscopy for 20 minutes (A) or 8 hours (B). First (i) and last (ii) timepoint of the time-lapse are shown, as well as the temporal color-coded image of the entire time-lapse (iii). CCSs that were present for a limited amount of time are shown in a single color from the color code (yellow arrows), while CCSs that were present for a longer time or persisted for the whole duration of the measurement are shown in the overlay color (white arrows). Scale bars 5 μm, except scale bars of enlargements 1 μm.

### 2. Divergent clathrin topology can be quantified in single-molecule localization microscopy

To accurately quantify the distribution of clathrin and avoid erroneous interpretation of CCSs located closely to each other, we visualized clathrin topology (mEos3.2-CLTA) using PALM-TIRF single-molecule localization microscopy (SMLM) (Fig 3i-ii). We selected TIRF illumination since it eliminates the background fluorescence signal from the cytoplasm, allowing for more precise visualization of membrane-associated protein organization (Fig S3). Consistent with our confocal results, we detected predominantly classical CCSs in the poorly metastatic KM12C cells, and a mix of classical CCSs and alternative FCLs in the metastatic KM12 cells (Fig 4Ai-iii and Fig S4i-iii). Of note, subtle discrepancies were also present within the same cell line (Fig S5A-B). To quantify the apparent differences between poorly and highly metastatic cells, we developed a two-step analysis workflow to identify and classify clathrin clusters as classical CCSs or alternative FCLs (Fig 3), leveraging the single-molecule information content of the SMLM imaging strategy.

**Figure 3:**
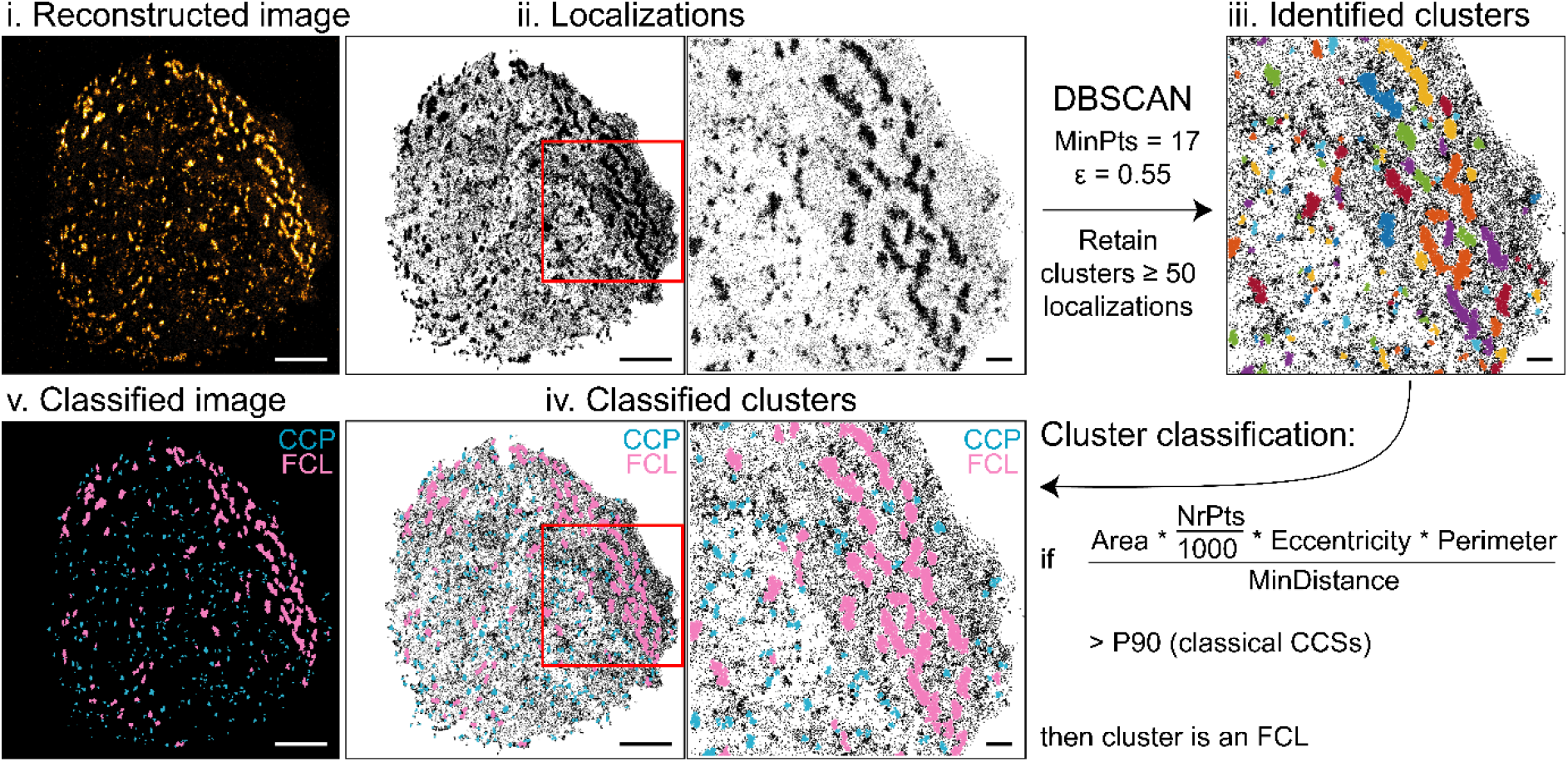
Cluster identification and classification workflow illustrated in a metastatic KM12L4a cell. Individual localizations (ii) that make up the PALM-TIRF image (i) were clustered using the DBSCAN algorithm with minimum 17 neighbors (MinPts) and search radius (ε) 0.55. Retaining clusters with ≥ 50 localizations resulted in identification of individual clathrin clusters (iii). The designed cluster classification model (iv - v) classifies these as either a classical CCS (blue, indicated as CCP) or an alternative FCL (pink) based on 5 cluster-specific parameters (cluster area, number of localizations NrPts, eccentricity, perimeter, and the distance to the nearest neighboring cluster MinDistance) and a global threshold. Scale bars 5 μm, except scale bars of enlargements 1 μm.

**Figure 4:**
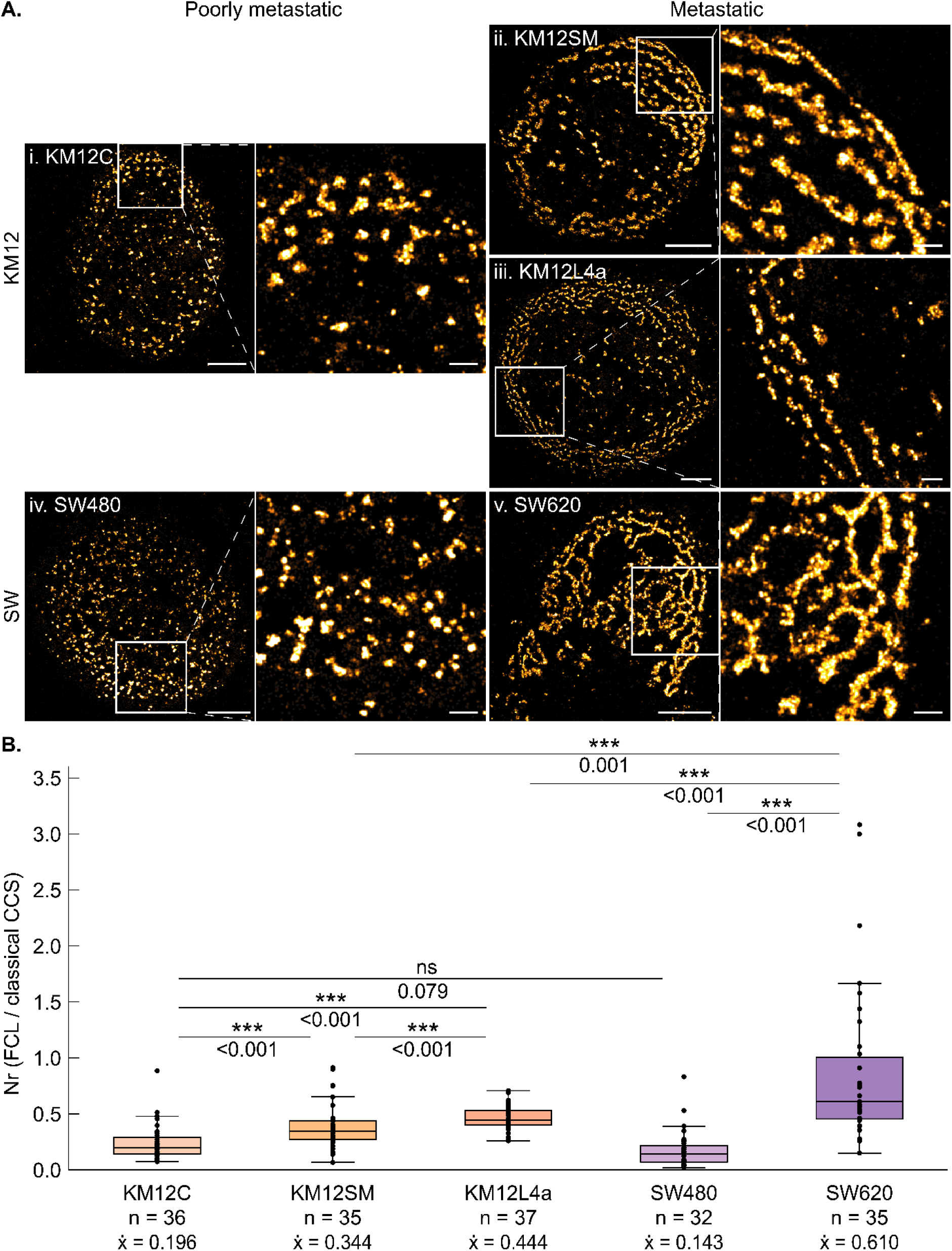
Visualization and quantification of clathrin topology at the ventral membrane of poorly and highly metastatic cells of the KM12 and SW models. A: Representative PALM-TIRF images of clathrin (mEos3.2-CLTA) at the ventral membrane of the different cell lines are shown. Scale bars 5 μm, except scale bars of enlargements 1 μm. B: Quantification of clathrin topology using the ratio of number of clusters classified as alternative FCLs versus classical CCSs per measured cell. Note that most cells have more classical CCSs than alternative FCLs (ratio < 1), except in 8 of the 35 SW620 cells. Data are plotted per cell line and the number of datapoints (n) and medians 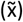 are provided. Statistical significances are shown on the plot (* significant difference for p ≤ 0.05; ** significant difference for p ≤ 0.01; *** significant difference for p ≤ 0.001; ns non-significant difference for p > 0.05).

For cluster identification, we applied DBSCAN density-based clustering that is capable of detecting clusters in a noisy dataset without making assumptions about their shape or size (34,35). After optimizing DBSCAN cluster parameters (MinPts and ε, Fig S6Aiii), and after discarding clusters with less than 50 localizations (corresponding to single clathrin molecules rather than a CCS, Fig S6B), the DBSCAN algorithm correctly identified individual clathrin clusters (Fig 3iii).

For cluster topology classification, different CCS characteristics such as cluster area, brightness and roundness have previously been applied (4,6,9,36,37). However, there seems to be a lack of unambiguous guidelines for parameter selection, so that different strategies rely on different CCS characteristics. Therefore, we designed a classification model that integrates multiple relevant CCS characteristics and considers – besides cluster area, brightness and roundness – the circumference and the distance to the nearest neighboring cluster. We found that these two additional parameters were effective in distinguishing between divergent CCSs (Fig S6C-D). One advantage of our integrated model is that it can rely on a global threshold for the combined parameters (Fig 3iv), rather than relying on threshold values for the individual parameters for which a consensus seems to be lacking in the literature. We set the global threshold based on an internal reference determined for 1247 classical and 1042 alternative CCSs that were manually selected, ensuring that maximally 10% of the classical CCSs are misclassified. Using this automated analysis workflow (see Materials and Methods section 8) we can quickly, accurately and objectively discriminate between classical and alternative clathrin topology in large SMLM datasets containing thousands of clusters (Fig 3iv-v).

To evaluate the effectiveness of our classification strategy, we examined the area of classified CCSs (Fig S6E), which is frequently used in the literature to identify alternative CCSs (4,6,9,36,37). We found that the area of classical CCSs is cell-type independent for the KM12 cell model with an average area of 33 000 nm^2^ (corresponding to a diameter of 205 nm when assuming a spherically shaped CCS). These results are in line with projections from the general CME model and with previous reports (4,6,9,37) considering that the resolution of optical microscopy yields slightly larger values compared to higher-resolution techniques like electron microscopy. In addition, we observed that alternative FCLs have an average area of 191 333 nm^2^, which is in line with the > 100 000 nm^2^ benchmark reported by Grove et al. (6) and is on average 5.79 times larger than a classical CCS. Unlike classical CCSs, FCL sizes vary significantly between the different cell lines, with larger FCLs associated to a higher metastatic potential. We also verified that cluster area is not correlated to cell area (Fig S6F). Overall, the similarities between our results and recent findings support our cluster classification model for the quantification of clathrin clusters in a SMLM dataset.

### 3. FCLs are associated to metastatic potential in two different CRC models, as quantified with single-molecule localization microscopy

Having verified our analysis strategy, we quantified clathrin topology in the KM12 model by comparing the number of clusters classified as alternative FCLs with those classified as classical CCSs, where this ratio was calculated for each of the measured cells (Fig 4B). We found that within a given KM12 cell, there are more classical CCSs than alternative FCLs since the ratio is lower than 1. Importantly, this ratio is significantly higher for metastatic KM12 cells than for their poorly metastatic counterpart (Fig 4B), which is due to a combination of significantly less classical CCSs and significantly more alternative FCLs (Fig S7). These results demonstrate that FCLs are associated with metastatic potential in the KM12 CRC model.

To reinforce our findings, we extended our research to the SW model for CRC metastasis. Clathrin topology imaged by PALM-TIRF microscopy is strikingly similar to that in the KM12 model, with predominantly small dot-like classical CCSs in the poorly metastatic SW480 cells, and a combination of classical CCSs and large, elongated FCLs in the metastatic SW620 cells (Fig 4Aiv-v and Fig S4iv-v). However, in the metastatic SW620 cells, clathrin seems to be reorganized even more extensively, since clathrin forms highly connected rosette-like structures that sometimes cover the entire ventral membrane with few classical CCSs present (Fig S5C). When applying our classification model on the SW dataset, we validated again that classical CCS area is cell-type independent (Fig S6E), and found that alternative FCLs are on average 5.82 times larger than classical CCSs. FCLs are significantly larger in the metastatic SW620 cell line compared to the poorly metastatic SW480 cell line (Fig S6E), and cluster area is again independent of cell area (Fig S6F). When quantifying clathrin topology in the SW model using the ratio of FCLs and classical CCSs (Fig 4B), we found that most cells have more classical CCSs than alternative FCLs. The poorly metastatic cells of both cell models are not significantly different (Fig 4B) and have a similar percentage classical CCSs and a similar percentage alternative FCLs (Fig S7). The metastatic SW620 cells have a significantly higher ratio when compared to their poorly metastatic counterpart (Fig 4B), and when compared to the metastatic cells of the KM12 model (Fig 4B), in accordance with the rosette-like structures observed in the SW620 cells. In conclusion, FCLs are associated with metastatic potential in both KM12 and SW CRC cell models.

Since protein expression and cellular function are closely related, we set out to investigate whether the aberrant clathrin topology could be due to altered expression levels of certain CME actors. We quantified the expression of clathrin light chain A (CLTA), clathrin heavy chain 1 (CLTC) and AP2 subunit α2 (AP2A2, an adaptor protein involved in CME) in the KM12 and SW models via Western blot (Fig S8). When comparing poorly and highly metastatic cells within the same cell model, we found that AP2A2 expression is downregulated in metastatic SW cells, while it is highly upregulated in metastatic KM12 cells, the latter as previously reported for this cell model (20). In terms of clathrin expression, we found that both CLTA and CLTC protein levels either remain unchanged or are downregulated in metastatic KM12 cells, while an opposite trend was seen within the SW model (CLTA is downregulated, but CLTC is upregulated in metastatic SW cells). Considering the differences between the KM12 and SW models, it is reasonable to assume that although alterations in CME actors might be inherent to CRCs’ metastatic progression, the exact changes may be associated to the different tropism of the cancer cells. It is noteworthy that clathrin protein expression does not seem to cause the aberrant clathrin topology observed in these cell lines, as one might intuitively expect. In addition, our results indicate that besides protein expression levels there must be additional factors influencing clathrin topology that converge to a similar phenotype albeit via different molecular mechanisms.

## Discussion

In this report, we show for the first time an association between metastatic potential and membranous clathrin distribution in two isogenic cell models representative of CRC metastasis. Confocal imaging revealed that poorly metastatic cells predominantly display classical dot-like CCSs, while metastatic cells exhibit both classical CCSs and larger, more elongated CCSs that we identified as FCLs based on their reported topology and lifetime. We observed FCLs for more than 8 hours in our measurements, an extremely extended lifetime that has to the best of our knowledge not yet been reported. When quantifying detailed clathrin topology using advanced fluorescence microscopy, we found that FCLs can be detected in both highly and poorly metastatic CRC cells, but that they are far more prevalent in cells with a higher metastatic potential.

While there is a clear link between metastatic potential and aberrant clathrin organization, the cause of FCL formation in these cell models remains an enigma since our analysis could not pinpoint the divergent topology to the expression of a specific CME component. It was previously reported that both *AP2M1* expression levels and alternative splicing of CLTC can influence clathrin organization (37,38), and therefore it could be that FCL formation cannot be attributed to one specific alteration. Instead, FCL formation may be cell type-specific or could depend on (interaction with) yet other proteins that were not analyzed here. For instance, there is growing evidence of an intimate relationship between FCLs and integrin αVβ5-containing reticular adhesions (14,39–41). Alternatively, mechanobiological cues could play a role in FCL formation, since an altered plasma membrane tension (12) and certain properties of the extracellular environment like rigidity and presentation of adhesion ligands (12,36,41–44) were reported to affect the presence of alternative CCSs.

Given the role of FCLs as a platform for sustained receptor signaling and their connection to cell proliferation (6,8,9,36), we hypothesize that alternative clathrin organization might be involved in cancer progression and metastasis. In addition, considering their close association with integrins and the extracellular matrix, these structures may be relevant to the migration of metastatic cancer cells during tumor dissemination. Further investigation of the link between metastatic potential and aberrant clathrin organization will contribute to unraveling the function of clathrin in tumor biology.

## Author contributions

C.C. and A.T. performed the optical imaging experiments and analyzed the data. Quantification of protein expression was performed by G.S.-F. and C.C.. R.N. assisted with the acquisition of images using the confocal microscope. S.R. and C.C. designed the experiments and wrote the final manuscript, with input from all the co-authors. J.H., R.B. and S.R. supervised the work.

## Acknowledgements

The authors would like to thank colleagues from KU Leuven MIP division, especially from the group of Prof. Rocha, for their input and critical questions. We thank Fidler’s lab (MD Anderson Cancer Center) for sharing KM12 model cell lines and Dr. Zhuang for making the EYFP-CLTA plasmid available (Addgene plasmid # 20921). We also thank Prof. R. Vitale (Université de Lille) for guidance and feedback on the statistical analysis. This work was funded by the Research Foundation - Flanders (C.C. is recipient of a PhD fellowship for fundamental research, FWO Grant Number 1121221N. G.S.-F. is recipient of a predoctoral contract, FWO Grant Number 1193818N), and the AES-ISCIII program to R.B. (PI17CIII/00045 and PI20CIII/00019 grants partially supported by FEDER funds). J.H. acknowledges financial support from the Research Foundation - Flanders (FWO Grant Numbers G0C1821N and ZW15 09-G0H6316N), from the Flemish government through long-term structural funding Methusalem (CASAS2, Meth/15/04), and from the MPI as a fellow. S.R. acknowledges financial support from KU Leuven (Grant Numbers KA/20/026 and IDN/20/021).

## Data availability

Data and analysis routines are available from the corresponding author upon request.

**Materials and methods** are available in the supplementary information.

## Notes

### Competing Interest Statement

The authors have declared no competing interest.

## References

1. Sochacki KA, Taraska JW. From flat to curved clathrin: controlling a plastic ratchet. Trends Cell Biol. 2019;29(3):241–56.

2. Heuser J, Evans L. Three-dimensional visualization of coated vesicle formation in fibroblasts. J Cell Biol. 1980;84(3):560–83.

3. Engqvist-Goldstein AEY, Zhang CX, Carreno S, Consuelo B, Heuser JE, Drubin DG. RNAi-mediated Hip1R silencing results in stable association between the endocytic machinery and the actin assembly machinery. Mol Biol Cell. 2004;15(4):1666–1679.

4. Merrifield CJ, Perrais D, Zenisek D. Coupling between clathrin-coated-pit invagination, cortactin recruitment, and membrane scission observed in live cells. Cell. 2005;121(4):593–606.

5. Saffarian S, Cocucci E, Kirchhausen T. Distinct dynamics of endocytic clathrin-coated pits and coated plaques. PLoS Biol. 2009;7(9):1–18.

6. Grove J, Metcalf DJ, Knight AE, Wavre-Shapton ST, Sun T, Protonotarios ED, et al. Flat clathrin lattices: stable features of the plasma membrane. Mol Biol Cell. 2014;25(22):3581–94.

7. Taylor MJ, Perrais D, Merrifield CJ. A high precision survey of the molecular dynamics of mammalian clathrin-mediated endocytosis. PLoS Biol. 2011;9(3):1–23.

8. Lampe M, Pierre F, Al-Sabah S, Krasel C, Merrifield CJ. Dual single-scission event analysis of constitutive transferrin receptor (TfR) endocytosis and ligand-triggered β2-adrenergic receptor (β2AR) or Mu-opioid receptor (MOR) endocytosis. Mol Biol Cell. 2014;25(19):3070–80.

9. Leyton-Puig D, Isogai T, Argenzio E, Van Den Broek B, Klarenbeek J, Janssen H, et al. Flat clathrin lattices are dynamic actin-controlled hubs for clathrin-mediated endocytosis and signalling of specific receptors. Nat Commun. 2017;8(16068):1–14.

10. Ehrlich M, Boll W, Van Oijen A, Hariharan R, Chandran K, Nibert ML, et al. Endocytosis by random initiation and stabilization of clathrin-coated pits. Cell. 2004;118(5):591–605.

11. Loerke D, Mettlen M, Yarar D, Jaqaman K, Jaqaman H, Danuser G, et al. Cargo and dynamin regulate clathrin-coated pit maturation. PLoS Biol. 2009;7(3):0628–39.

12. Baschieri F, Porshneva K, Montagnac G. Frustrated clathrin-mediated endocytosis – causes and possible functions. J Cell Sci. 2020;133(11):1–10.

13. Lampe M, Vassilopoulos S, Merrifield C. Clathrin coated pits, plaques and adhesion. J Struct Biol. 2016;196(1):48–56.

14. Zuidema A, Wang W, Kreft M, Bleijerveld OB, Hoekman L, Aretz J, et al. Molecular determinants of αVβ5 localization in flat clathrin lattices – role of αVβ5 in cell adhesion and proliferation. J Cell Sci. 2022;135(11):1–18.

15. Ramsay AG, Marshall JF, Hart IR. Integrin trafficking and its role in cancer metastasis. Cancer Metastasis Rev. 2007;26(3-4):567–78.

16. Lanzetti L, Di Fiore PP. Endocytosis and cancer: an “insider” network with dangerous liaisons. Traffic. 2008;9(12):2011–21.

17. Mosesson Y, Mills GB, Yarden Y. Derailed endocytosis: an emerging feature of cancer. Nat Rev Cancer. 2008;8(11):835–50.

18. Elkin SR, Bendris N, Reis CR, Zhou Y, Xie Y, Huffman KE, et al. A systematic analysis reveals heterogeneous changes in the endocytic activities of cancer cells. Cancer Res. 2015;75(21):4640–50.

19. Khan I, Steeg PS. Endocytosis: a pivotal pathway for regulating metastasis. Br J Cancer. 2020;124(1):66–75.

20. Mendes M, Peláez-García A, López-Lucendo M, Bartolomé RA, Calviño E, Barderas R, et al. Mapping the spatial proteome of metastatic cells in colorectal cancer. Proteomics. 2017;17(19):1–11.

21. Solís-Fernández G, Montero-Calle A, Martínez-Useros J, López-Janeiro Á, Ríos VDL, Sanz R, et al. Spatial Proteomic Analysis of Isogenic Metastatic Colorectal Cancer Cells Reveals Key Dysregulated Proteins Associated with Lymph Node, Liver, and Lung Metastasis. Cells. 2022;11(447):1–23.

22. Morikawa K, Walker SM, Jessup JM, Fidler I. In vivo selection of highly metastatic cells from surgical specimens of different primary human colon carcinomas implanted into nude mice. Cancer Res. 1988;48(7):1943–8.

23. Morikawa K, Walker SM, Nakajima M, Pathak S, Jessup JM, Fidler I. Influence of organ environment on the growth, selection, and metastasis of human colon carcinoma cells in nude mice. Cancer Res. 1988;48(23):6863–71.

24. Leibovitz A, Stinson JC, McCombs WB, McCoy CE, Mazur KC, Mabry ND. Classification of Human Colorectal Adenocarcinoma Cell Lines. Cancer Res. 1976;36(12):4562–9.

25. Hewitt RE, McMarlin A, Kleiner D, Wersto R, Martin P, Tsoskas M, et al. Validation of a model of colon cancer progression. J Pathol. 2000;192(4):446–54.

26. Hegde P, Qi R, Gaspard R, Abernathy K, Dharap S, Earle-Hughes J, et al. Identification of tumor markers in models of human colorectal cancer using a 19,200-element complementary DNA microarray. Cancer Res. 2001;61(21):7792–7.

27. Kuniyasu H, Ohmori H, Sasaki T, Sasahira T, Yoshida K, Kitadai Y, et al. Production of Interleukin 15 by Human Colon Cancer Cells Is Associated with Induction of Mucosal Hyperplasia, Angiogenesis, and Metastasis. Clin Cancer Res. 2003;9(13):4802–10.

28. Li A, Varney ML, Singh RK. Constitutive expression of growth regulated oncogene (gro) in human colon carcinoma cells with different metastatic potential and its role in regulating their metastatic phenotype. Clin Exp Metastasis. 2005;21(7):571–9.

29. Barderas R, Bartolomé RA, Fernandez-Aceñero MJ, Torres S, Casal JI. High expression of IL-13 receptor α2 in colorectal cancer is associated with invasion, liver metastasis, and poor prognosis. Cancer Res. 2012;72(11):2780–90.

30. Calon A, Espinet E, Palomo-Ponce S, Tauriello DVF, Iglesias M, Céspedes MV, et al. Dependency of Colorectal Cancer on a TGF-β-Driven Program in Stromal Cells for Metastasis Initiation. Cancer Cell. 2012;22(5):571–84.

31. Solís-Fernández G, Montero-Calle A, Sánchez-Martínez M, Peláez-García A, Fernández-Aceñero MJ, Pallarés P, et al. Aryl hydrocarbon receptor-interacting protein regulates tumorigenic and metastatic properties of colorectal cancer cells driving liver metastasis. Br J Cancer. 2022;136:1604–15.

32. Montero-Calle A, Gómez de Cedrón M, Quijada-Freire A, Solís-Fernández G, López-Alonso V, Espinosa-Salinas I, et al. Metabolic Reprogramming Helps to Define Different Metastatic Tropisms in Colorectal Cancer. Front Oncol. 2022;12(July):1–19.

33. Gaidarov I, Santini F, Warren RA, Keen JH. Spatial control of coated-pit dynamics in living cells. Nat Cell Biol. 1999;1(1):1–7.

34. Ester M, Kriegel H-P, Sander J, Xu X. A density-based algorithm for discovering clusters in large spatial databases with noise. KDD-96 Proc. 1996;226–31.

35. Khater IM, Nabi IR, Hamarneh G. A Review of Super-Resolution Single-Molecule Localization Microscopy Cluster Analysis and Quantification Methods. Patterns. 2020;1(3):100038.

36. Baschieri F, Dayot S, Elkhatib N, Ly N, Capmany A, Schauer K, et al. Frustrated endocytosis controls contractility-independent mechanotransduction at clathrin-coated structures. Nat Commun. 2018;9(3825):1–13.

37. Moulay G, Lainé J, Lemaitre M, Nakamori M, Nishino I, Caillol G, et al. Alternative splicing of clathrin heavy chain contributes to the switch from coated pits to plaques. J Cell Biol. 2020;219(9):1–16.

38. Dambournet D, Sochacki KA, Cheng AT, Akamatsu M, Taraska JW, Hockemeyer D, et al. Genome-edited human stem cells expressing fluorescently labeled endocytic markers allow quantitative analysis of clathrin-mediated endocytosis during differentiation. J Cell Biol. 2018;217(9):3301–11.

39. Lock JG, Jones MC, Askari JA, Gong X, Oddone A, Olofsson H, et al. Reticular adhesions are a distinct class of cell-matrix adhesions that mediate attachment during mitosis. Nat Cell Biol. 2018;20(11):1290–302.

40. Lock JG, Baschieri F, Jones MC, Humphries JD, Montagnac G, Strömblad S, et al. Clathrin-containing adhesion complexes. J Cell Biol. 2019;218(7):2086–95.

41. Hakanpää L, Abouelezz A, Lenaerts A, Algie M, Bärlund J, Katajisto P, et al. Flat clathrin lattices nucleate reticular adhesions in an integrin β1 activity-dependent manner. BioRxiv Prepr. 2022;1–13.

42. Elkhatib N, Bresteau E, Baschieri F, Rioja AL, Van Niel G, Vassilopoulos S, et al. Tubular clathrin/AP-2 lattices pinch collagen fibers to support 3D cell migration. Science (80-). 2017;356(6343).

43. Zuidema A, Wang W, Kreft M, Te Molder L, Hoekman L, Bleijerveld OB, et al. Mechanisms of integrin αVβ5 clustering in flat clathrin lattices. J Cell Sci. 2018;131(21):1–16.

44. Bresteau E, Elkhatib N, Baschieri F, Bellec K, Guérin M, Montagnac G. Clathrin-coated structures support 3D directed migration through local force transmission. Sci Adv. 2021;7(45):1–11.

