## Supplementary information for "Flat clathrin lattices are linked to metastatic potential in colorectal cancer"

Supplementary Figures and Tables………………………………………………………………………………………….. p. 2-10

Materials and Methods……………………………………………………………………………………………………… p. 11-15

Supplementary Info References…………………………………………………………………………………………… p. 15

### Supplementary Figures and Tables

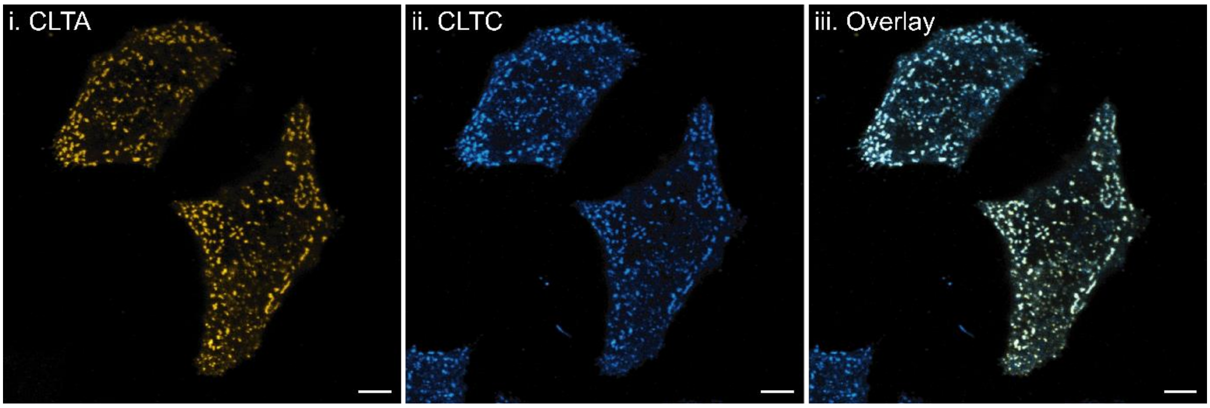


Figure S1: Clathrin topology imaged with dual-color confocal microscopy. Representative images of transiently transfected EYFP-CLTA (i) and immunolabeled Atto647N-CLTC (ii) at the ventral membrane of metastatic KM12L4a cells. Images were taken sequentially for the same field of view. Overlay (iii) shows a very comparable clathrin topology between transfection and immunolabeling conditions, except for the cell in the lower left corner that was not transfected and therefore only displayed CLTC. Scale bars 5 µm.

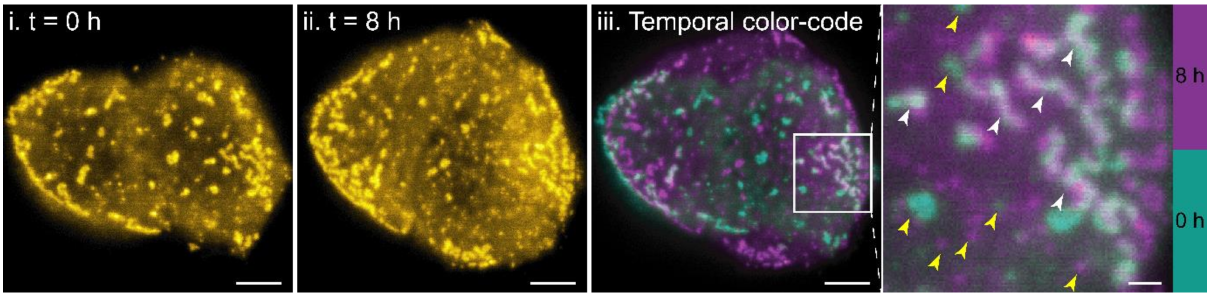


Figure S2: Clathrin dynamics at the ventral membrane of a metastatic KM12L4a cell transiently transfected with EYFP-CLTA, imaged by time-lapse confocal microscopy for 8 hours. First (i) and last (ii) timepoint of the time-lapse are shown, as well as the temporal color-coded image of the entire time-lapse (iii). CCSs that were present for a limited amount of time are shown in a single color from the color code (yellow arrows), while CCSs that were present for a longer time or persisted for the whole duration of the measurement are shown in the overlay color (white arrows). Scale bars 5 µm, except scale bar of the enlargement 1 µm.

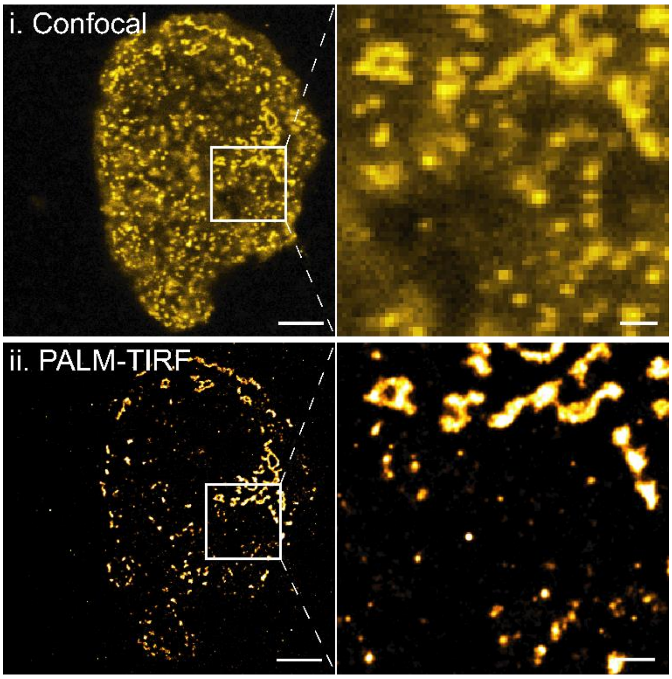


Figure S3: Comparison of confocal and PALM-TIRF microscopy for clathrin imaging at the ventral membrane. The same metastatic KM12L4a cell transfected with mEos3.2-CLTA was imaged using confocal microscopy (i) and PALM-TIRF microscopy (ii). Note that CCSs in the cytoplasm are visible (as out-of-focus structures) in the confocal image, while the PALM-TIRF image contains only CCSs that are truly located at the ventral membrane due to the enhanced axial resolution in TIRF-mode. Scale bars 5 µm, except scale bars of enlargements 1 µm.

i. KM12C

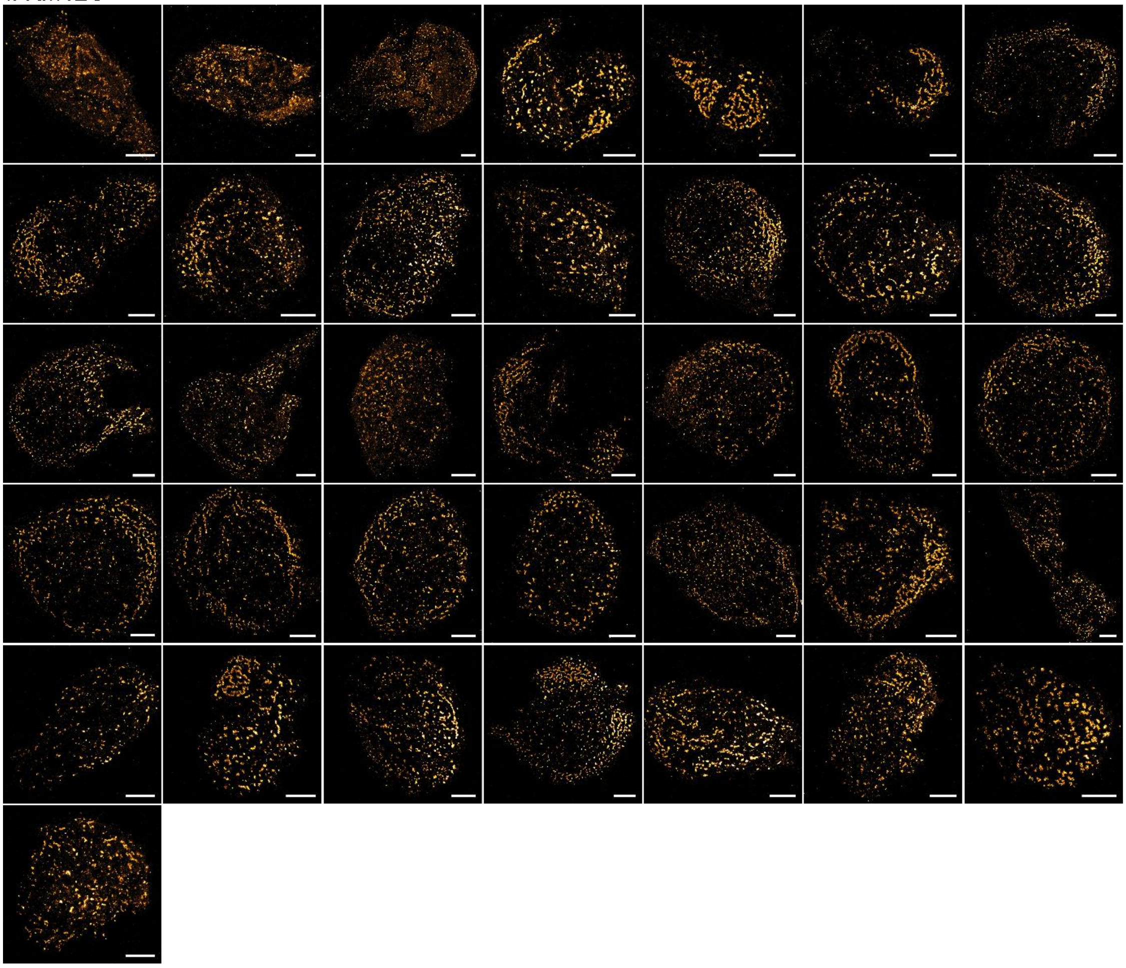

ii. KM12SM

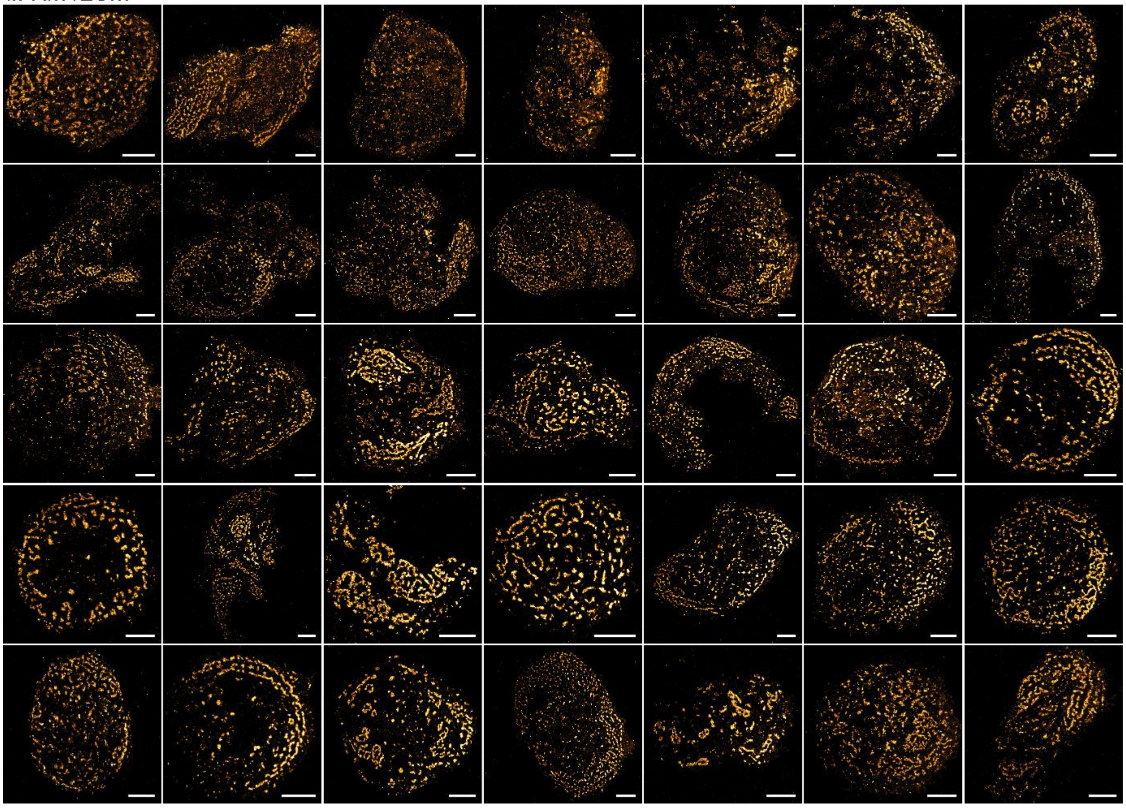

iii. KM12L4a

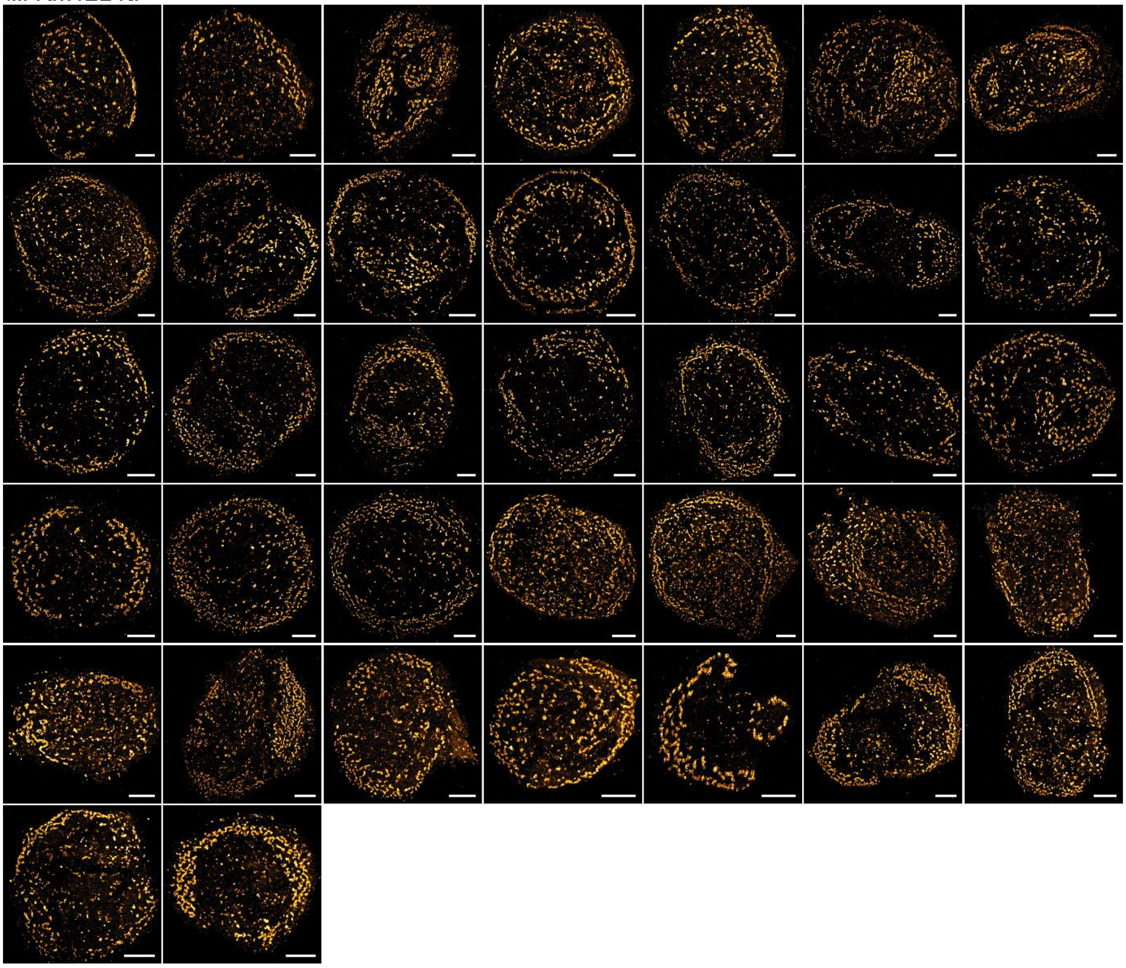

iv. SW480

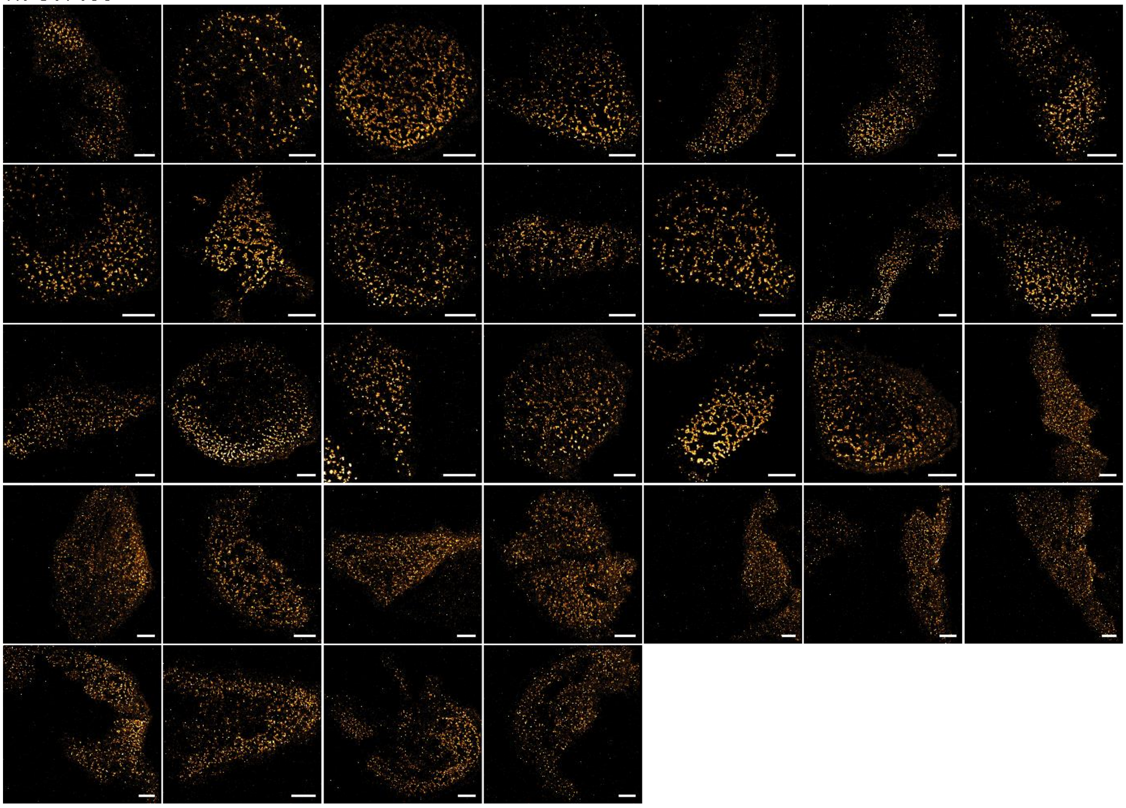

v. SW620

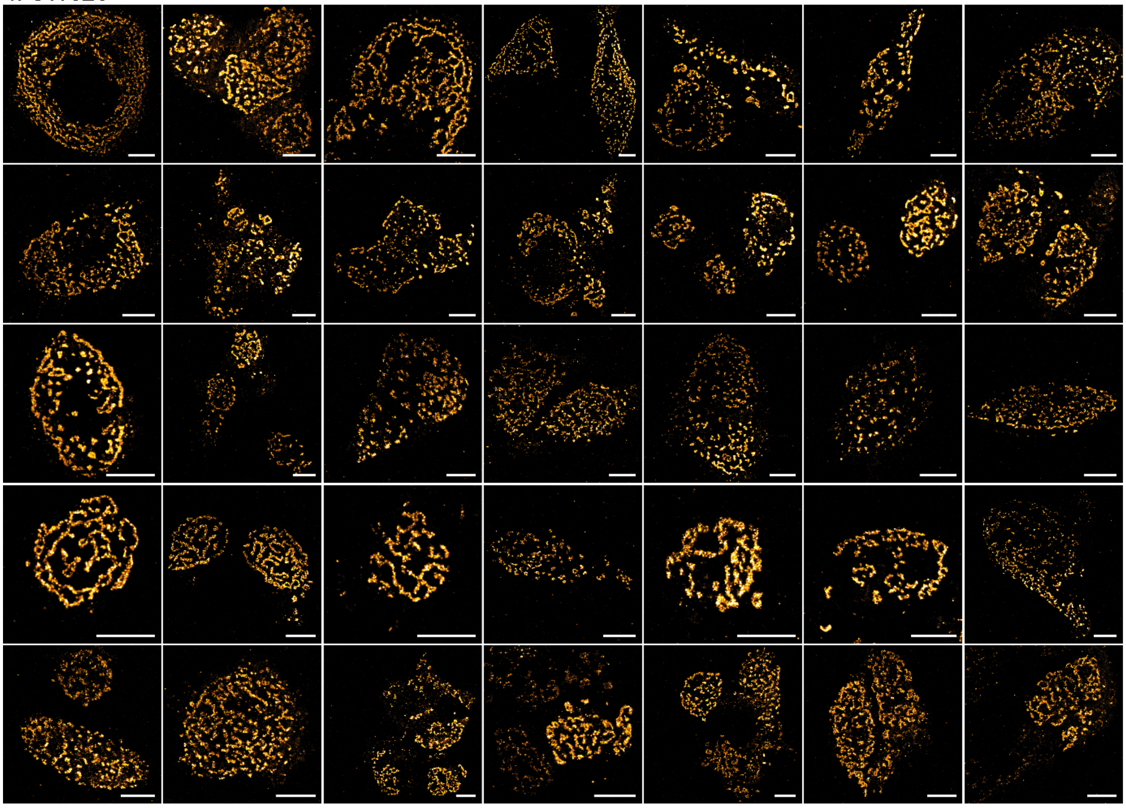

Figure S4: Complete PALM-TIRF dataset of clathrin (mEos3.2-CLTA) imaged at the ventral membrane of KM12 (i-iii) and SW (iv-v) cells. Minimally 32 cells from at least 7 biological replicates were measured per cell line. Images are enlargements of the measured field of view to only contain the cell that was isolated for the analysis where possible. Scale bars 5 µm.

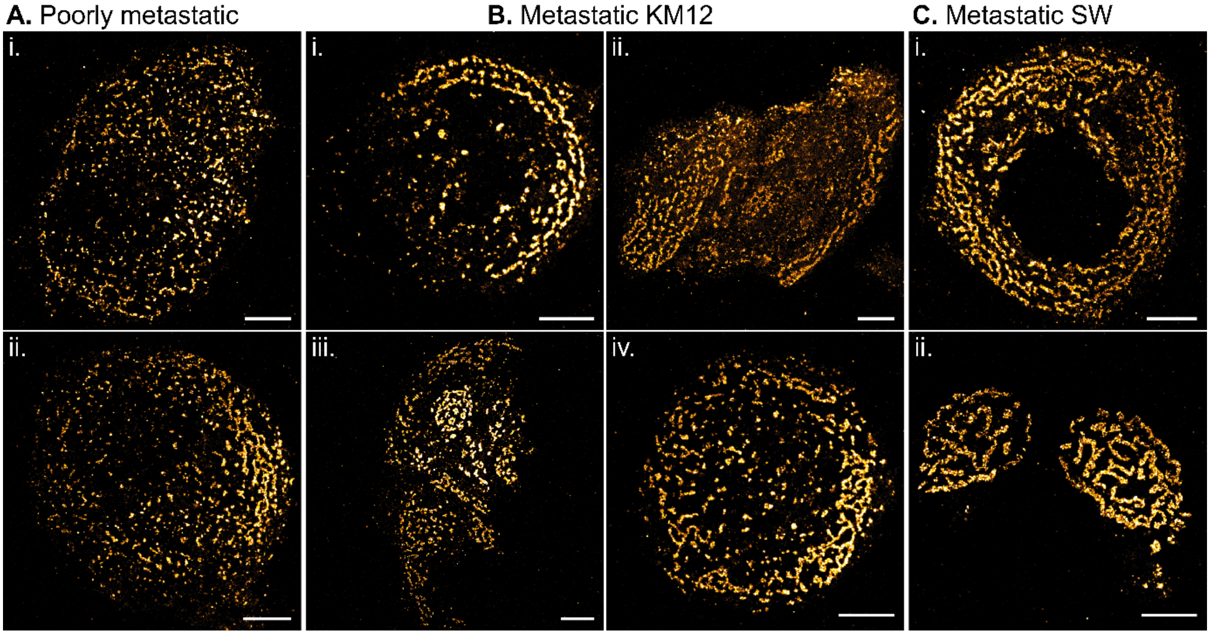


Figure S5: Variability of clathrin topology (mEos3.2-CLTA) at the ventral membrane of the KM12 and SW models, imaged with PALM-TIRF microscopy. Images representative of the variability found within poorly metastatic cells (A), within metastatic KM12 cells (B) and within metastatic SW cells (C) are shown. Poorly metastatic cells predominantly display classical CCSs (Ai), however, occasionally certain membrane regions also exhibit alternative FCLs (Aii), but to a far less extent than in the metastatic cells. Especially in the metastatic cells lines there is a larger discrepancy in clathrin distribution since FCLs in the metastatic KM12 cells can be arranged in concentric rings following the cell periphery (Bi), in patterns in certain peripheral parts (Bii), in patches more in the central part (Biii), or as FCLs with a higher degree of connectivity (Biv). The metastatic SW620 cells generally have many FCLs that are organized as patterned FCLs (Ci), or as highly connected rosette-like structures that cover a large portion of the ventral membrane (Cii). Scale bars 5 µm.

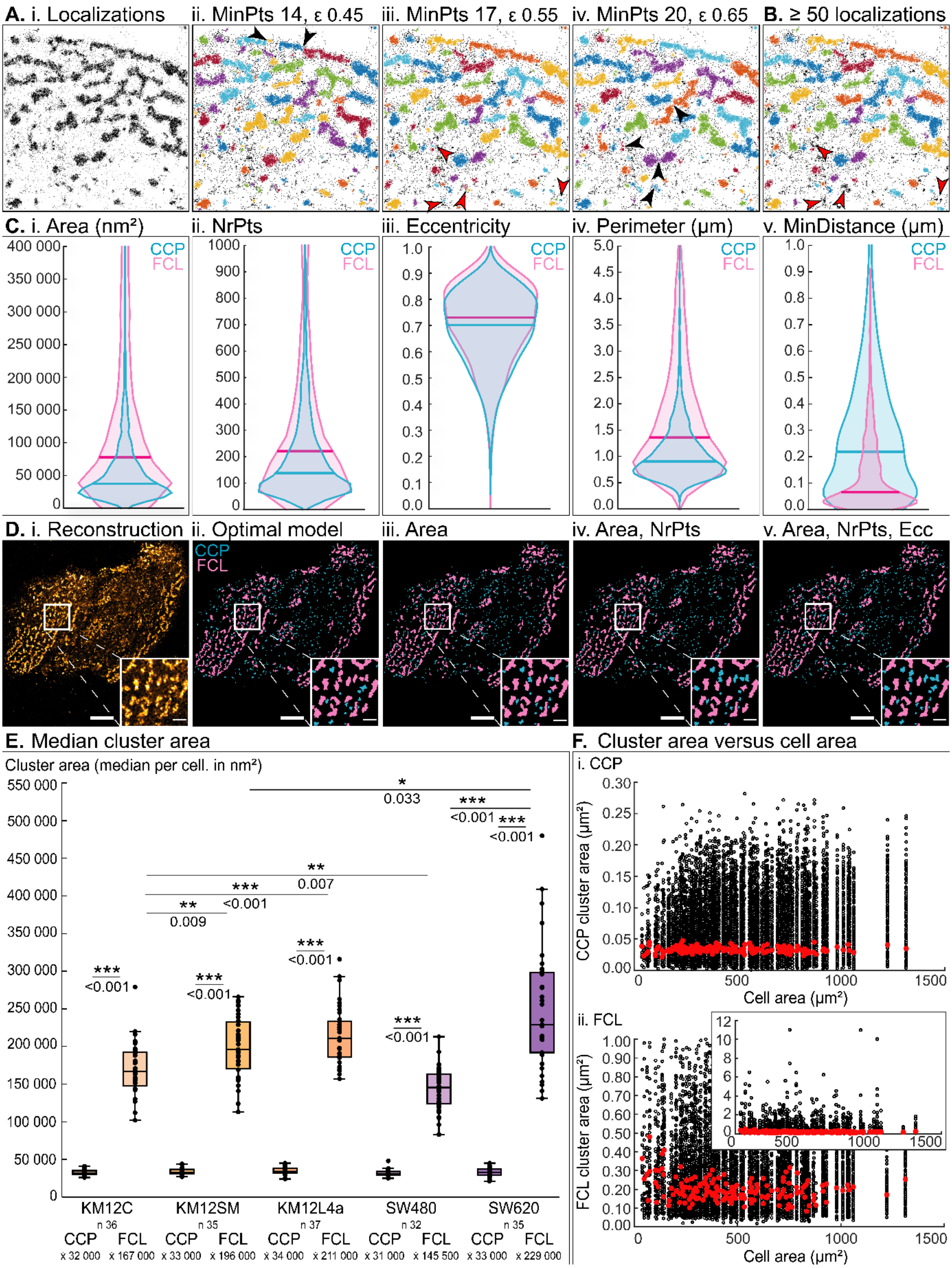


Figure S6: Optimization and testing of the designed cluster classification model.

A: Optimization of DBSCAN cluster parameters where minimum number of neighbors (MinPts) and search radius (ε) are optimized (Aii-iv) based on visual observation of a cellular region (Ai). Note that all combinations of MinPts = 8 – 23 and ε = 0.35 – 0.65 were attempted, but that only three parameter combinations are illustrated here. Black arrows indicate examples of suboptimal clustering, identifying MinPts = 17 and ε = 0.55 (Aiii) as optimal DBSCAN cluster parameters.

B: Justification of the ≥ 50 localization criterium. Clusters that were identified by DBSCAN (Aiii) but have < 50 localizations are not retained for further analysis (examples indicated by red arrows in B and Aiii for comparison). These clusters are extremely small, and their omission does not change our global view of true CCSs.

C: Distribution of cluster characteristics for classical CCSs (blue, indicated as CCP) and alternative FCLs (pink). Cluster characteristics were cluster area (i), the number of localizations within the cluster (ii, NrPts, associated with cluster brightness), the cluster eccentricity (associated with cluster roundness), the cluster perimeter and the distance to the nearest neighboring cluster (MinDistance). These characteristics have a different distribution for classical CCSs and alternative FCLs, and were hence employed in the cluster classification model. Note that FCL distributions and medians are shifted towards higher values compared to classical CCS, except for MinDistance. These relative shifts were accounted for when constructing the classification model (i.e. MinDistance was placed in the denominator of the classification model). Plotted values are derived from 1247 classical CCSs and 1042 alternative FCLs that we manually selected from 3-5 different cellular regions containing predominantly classical CCSs and alternative FCLs, respectively. Violin plots were made using the open-source MATLAB Violin plot function (1).

D: Illustration of the performance of cluster perimeter and MinDistance (ii) compared to when only considering cluster area (iii); cluster area and NrPts (iv); or cluster area, NrPts and eccentricity (v). Global thresholds for each iteration were determined as performed for the optimized classification model, namely so that maximally 10% of the classical CCSs were misclassified. Scale bars 5 µm, except scale bars of enlargements 1 µm.

E: Median cluster area of classified CCSs per measured cell. Data are plotted per classified category (CCP for classical CCSs, or FCL) and per cell line. The number of datapoints (n) and medians (x̃) are provided. Significant differences are calculated for CCP and FCL categories of the same cell line, for relevant CCP categories of different cell lines and for relevant FCL categories of different cell lines. These relevant categories of different cell lines are defined as: KM12C-KM12SM, KM12C-KM12L4a, KM12SM-KM12L4a, SW480-SW620, KM12C-SW480, KM12SM-SW620, KM12L4a-SW620. Statistical significances are shown (* significant difference for $p \leq 0.05$; ** significant difference for $p \leq 0.01$; *** significant difference for $p \leq 0.001$), while non-significant differences ($p > 0.05$) are not displayed.

F: Area of individual CCSs plotted in function of cell area. Separate plots are made for the identified classical CCSs (i) and alternative FCLs (ii), where medians are shown as red dots. Graphs contain PALM-TIRF data from KM12 and SW models, where data is plotted with increasing cell area. While extremely small cells seem to present smaller classical CCSs, there is no significant relationship between classical CCS area and cell area, or between FCL area and cell area. This implies that cluster area does not depend on cell area. Inset for alternative FCLs (ii) shows the true dimensions of the data.

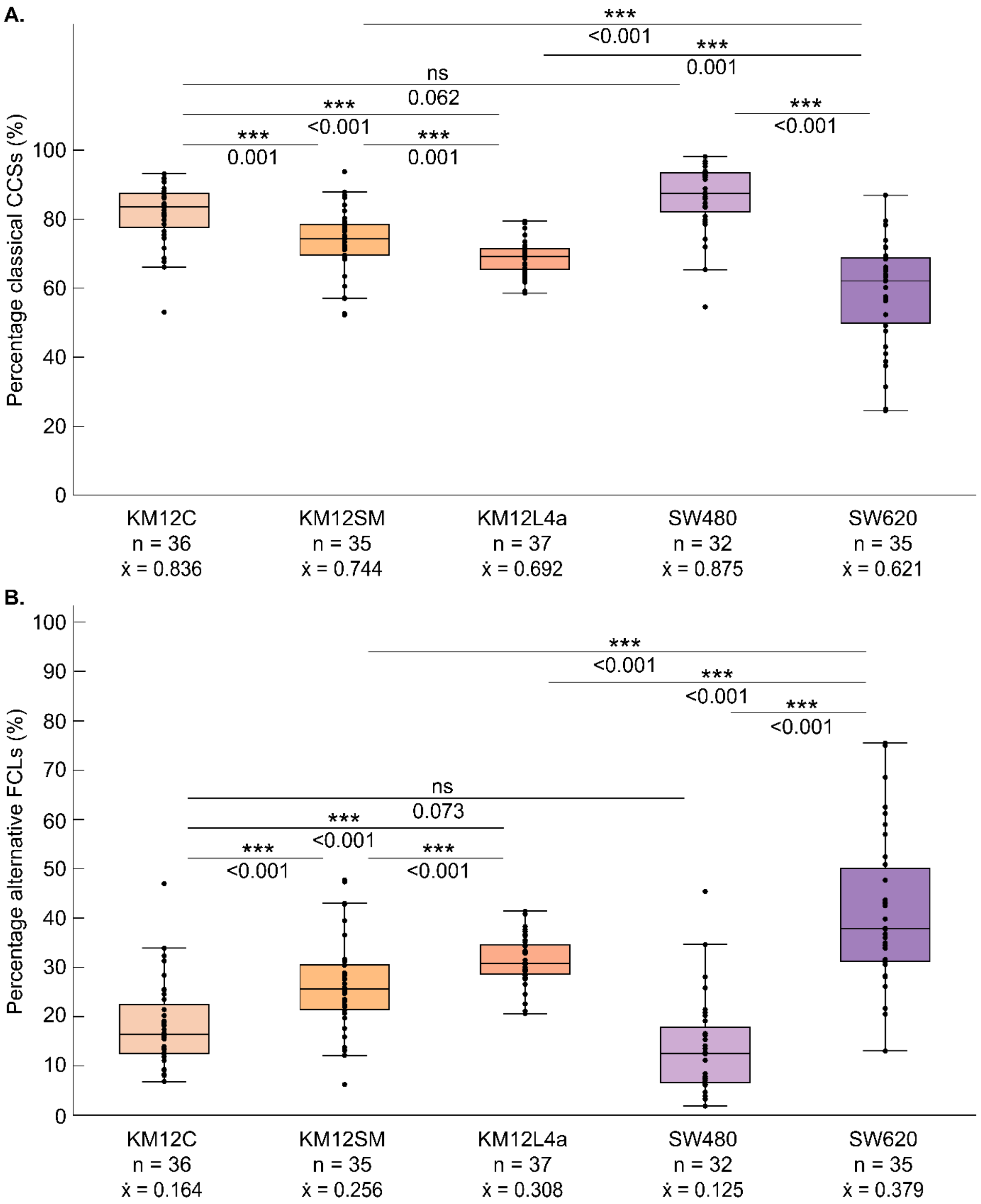


Figure S7: Percentage classical CCSs (A) or alternative FCLs (B) per measured cell. Percentages represent the number of CCS identified in each category, divided by the total number of identified clusters. Data are plotted per cell line and the number of datapoints (n) and medians (x̃) are provided. Statistical significances are shown on the plot (* significant difference for $p \leq 0.05$; ** significant difference for $p \leq 0.01$; *** significant difference for $p \leq 0.001$; ns non-significant difference for $p > 0.05$).

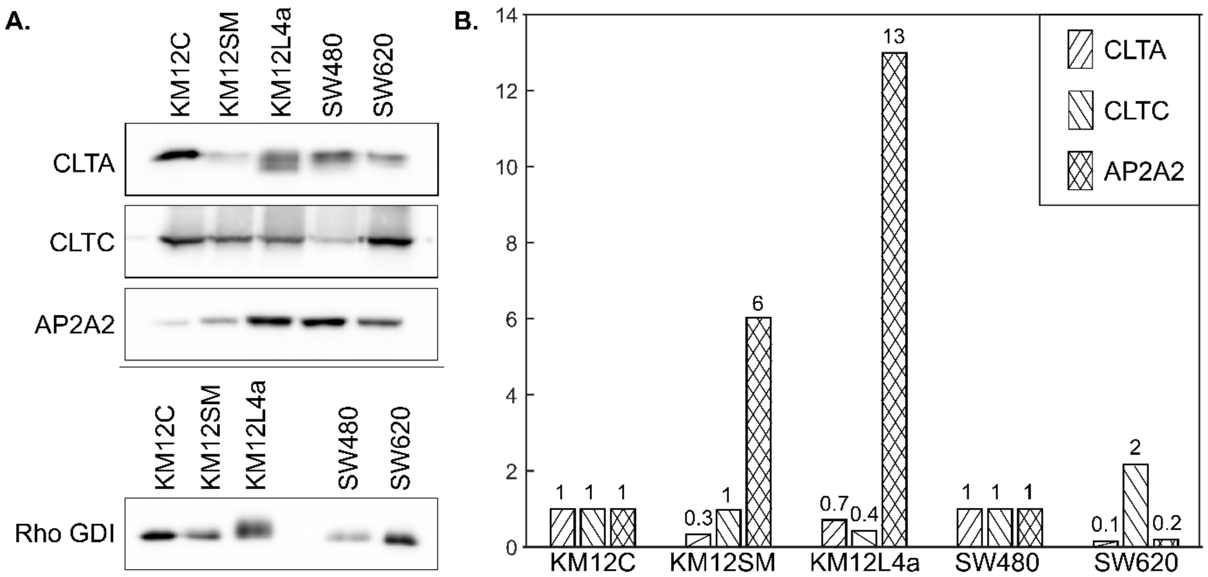


Figure S8: Expression levels of several CME actors. Western blot visualization (A) of the different target proteins and Rho GDI which served as loading control. Signal development was performed with exposure conditions optimized for each target. Rho GDI results were obtained from a second gel with the same amount of protein extracts loaded. Quantified normalized expression levels (B) were referenced to the poorly metastatic cell line of the corresponding cell model (KM12C for the KM12 model and SW480 for the SW model).

Table S1: Employed reagents for fluorescent labeling of targets in the KM12 and SW cell models.

| | **target** | **primary antibody** | **secondary antibody** |
|---|---|---|---|
| **Western blot** | clathrin light chain A (CLTA)<br>in KM12 or SW cells | rabbit anti-CLTA<br>(Proteintech, 10852-1-AP)<br>1/1000 dilution | Goat anti-rabbit (GAR)-HRP<br>conjugate<br>(Sigma, A6154)<br>1/2000 dilution |
| | clathrin heavy chain 1 (CLTC)<br>in KM12 or SW cells | mouse anti-CLTC<br>(Proteintech, 66487-1-Ig)<br>1/1000 dilution | Goat anti-mouse (GAM)-HRP<br>conjugate<br>(Sigma, A4416)<br>1/2000 dilution |
| | adaptor protein 2, α2-subunit (AP2A2)<br>in KM12 or SW cells | mouse anti-AP2A2<br>(Santa cruz, sc-55497)<br>1/1000 dilution | Goat anti-mouse (GAM)-HRP<br>conjugate<br>(Sigma, A4416)<br>1/2000 dilution |
| | Rho GDI<br>(protein normalization control)<br>in KM12 or SW cells | mouse anti-Rho GDI<br>(Santa cruz, sc-373724)<br>1/1000 dilution | Goat anti-mouse (GAM)-HRP<br>conjugate<br>(Sigma, A4416)<br>1/2000 dilution |
| | **target** | **plasmid construct** | **transfection conditions** |
| **Transient transfection** | clathrin light chain A (CLTA)<br>in KM12 or SW cells | EYFP-CLTA<br>(Addgene, 220921) | 0.9 µL TransIT-X2®<br>(Mirus, 6000)<br>0.3 µg plasmid DNA<br>~ 0.80 *$10^6$ cells<br>measurement after ~ 18 hours |
| | clathrin light chain A (CLTA)<br>in KM12 cells | mEos3.2-CLTA<br>(derived from EYFP-CLTA) | 3 µL FuGENE® 6<br>(Promega, E2691)<br>0.5 µg plasmid DNA<br>~ 0.55 * $10^6$ cells<br>measurement after ~ 48 hours |
| | clathrin light chain A (CLTA)<br>in SW cells | mEos3.2-CLTA<br>(derived from EYFP-CLTA) | 4.5 µL FuGENE® 6<br>(Promega, E2691)<br>0.75 µg plasmid DNA<br>~ 0.30 * $10^6$ cells<br>measurement after ~ 48 hours |
| | **target** | **primary antibody** | **secondary antibody** |
| **Immuno-fluorescence** | clathrin heavy chain 1 (CLTC)<br>in KM12 or SW cells | mouse anti-CLTC<br>(NovusBio, NB300-613)<br>1/500 dilution | Goat anti-mouse (GAM)-Atto647N<br>(Sigma, 50185)<br>1/1000 dilution |

**Materials and Methods**

1. Colorectal cancer cell culture

Isogenic KM12C, KM12SM and KM12L4a cell lines from Fidler's lab (MD Anderson Cancer Center) and isogenic SW480 and SW620 cell lines from ATCC (American Type Culture Collection) were used (2,3). Cell lines were maintained in a humidified incubator at 37 °C and 5% $CO_2$ in Dulbecco's Modified Eagle Medium (DMEM, Gibco, 31053028) supplemented with 10% Fetal Bovine Serum (FBS, Sigma, F7524), 1% GlutaMAX (Gibco, 35050038) and 0.1% Gentamycin (Carl Roth, 2475.1).

2. Western blot and quantification

Cells grown to 90% confluence were harvested using 1x Trypsin-EDTA (Thermo, 59418C), pelleted by centrifugation (5 min at 220 G) and the dry pellet was frozen at -20 °C for at least 1 h. Protein extracts were obtained by cell lysis using RIPA buffer mixed with phosphatases and proteases inhibitor cocktails (RIPA Sigma, R0278 with 1/100 phosphatase inhibitor MCE, HY-K0022 and 1/100 protease inhibitor MCE, HY-K0010) and 10 min centrifugation at 21 000 G. Next, 5 µg protein extracts quantified using the Tryptophan fluorescence method (4,5) were separated on a 10% SDS-PAGE gel under reducing conditions and transferred onto a nitrocellulose membrane for 90 min at 100 V. Protein transfer was verified by Ponceau staining (Sigma, P3504) which was afterwards removed by washes with DPBST (DPBS Thermo, 14200067 + 0.1% Tween20 Sigma, P1379). For specific target detection, the membrane was cut according to the expected band sizes, blocked in blocking buffer (3% skimmed milk in DPBST) for 1 h at room temperature, and incubated with primary antibody diluted in blocking buffer overnight at 4 °C. The membrane was washed 3x with DPBST for 10 min and incubated for 1 h at room temperature with secondary antibody diluted in blocking buffer. Washes were performed as before, chemiluminescent signal was generated with the ECL Pico PLUS Chemiluminescent Substrate (Thermo, 34580) and protein bands were imaged on an Amersham Imager 680 (GE Healthcare). Details of the antibodies used are listed in Table S1. For quantification, standard densitometry was performed using the ImageJ gel analysis tool (6,7). The peak area was measured in the lane plot and numerical values were further processed in Excel, normalized using Rho GDI signal as loading control, and referenced to the poorly metastatic cell line of the corresponding cell model for their comparison.

3. Plasmids and transfection

Plasmids for transient expression of clathrin light chain A (CLTA) were EYFP-CLTA (generated by Xiaowei Zhuang, Addgene plasmid EYFP-Clathrin, # 20921) (8), and mEos3.2-CLTA, generated by replacing the EYFP coding sequence by mEos3.2 using standard molecular cloning techniques. Reverse transfections were performed immediately after seeding the appropriate number of cells (Table S1) in a 35 mm disk with 14 mm glass-bottom insert (Cellvis, D35-14-1.5-N), a glass-bottom 6-well plate (Cellvis, P06-1.5H-N) or a 35 mm disk with imprinted grid (Ibidi, 80156). The transfection mixture for EYFP-CLTA was prepared using TransIT-X2® (Mirus, 6000) according to the Universal Transfection Reagent Protocol by Sigma-Aldrich (Sigma, T0956) with optimized reagent quantities (Table S1). Briefly, plasmid DNA was added to 50 µL DMEM with 1% GlutaMAX in a first tube (A), while TransIT-X2® was added to 50 µL DMEM with 1% GlutaMAX in a second tube (B). Both tubes were gently vortexed, tube B was immediately added to tube A, gently vortexed and spun down. The transfection mixture for mEos3.2-CLTA was prepared by first adding FuGENE® 6 (Promega, E2691) and then plasmid DNA to 100 µL DMEM with 1% GlutaMAX. For both methods the transfection mixture was added dropwise to recently seeded cells after 15-20 min incubation at room temperature, and the sample was incubated in a humidified incubator at 37 °C and 5% $CO_2$ for the appropriate time (Table S1).

### 4. Immunolabeling

Immunolabeling of clathrin heavy chain 1 (CLTC) was performed on EYFP-CLTA transfected samples in a 35 mm disk. Samples were washed 1x with 1 mL prewarmed DPBS (Thermo, 14200067), fixed for 10 min in 200 µL prewarmed PFA 4% (Thermo, 28908) in DPBS, then washed 3x with 1 mL DPBST (DPBS + 0.1% Tween20 Sigma, P1379) and permeabilized with 200 µL 0.1% Triton-X100 (Sigma, 9002-93-1) in DPBST for 15 min at room temperature. Three washes with 1 mL blocking buffer (10% FBS, Sigma, F7524 in DPBST) were performed and the sample was blocked for 1 h at room temperature. Next, the sample was incubated with 200 µL primary antibody diluted in blocking buffer overnight at 4 °C. Three washes with 1 mL blocking buffer were performed for 5 min each and an additional blocking step with 200 µL 5% goat serum (Sigma, G9023) in blocking buffer was performed for 15 min at room temperature. The sample was incubated with 200 µL Atto647N secondary antibody diluted in 5% goat serum blocking buffer for 1 h at room temperature in the dark. Two washes with 1 mL blocking buffer and two washes with 1 mL DPBST were performed for 7 min each before the sample was placed in DPBST and imaged. Details of the antibodies used are listed in Table S1.

### 5. Confocal fluorescence microscopy

Confocal imaging was performed on a Leica TCS SP8 microscope (Leica Microsystems GmbH) with 63x oil (NA: 1.4) or 63x water (NA: 1.2) immersion objective, for imaging clathrin topology and clathrin dynamics, respectively. Sample excitation was done using a supercontinuum laser (WLL, NKT Photonics) or a diode laser at 488 nm for mEos3.2, at 514 nm for EYFP or at 638 nm for Atto647N. Emission was captured on highly sensitive hybrid detectors (HyD SMD, Leica Microsystems GmbH) between 493-744 nm for mEos3.2-labeled samples, between 519-769 nm for EYFP-labeled samples, or between 519-638 nm and 643-779 nm for dual-color EYFP -and Atto647N samples, respectively. Transmission images were captured simultaneously on a transmitted light detector. Image acquisition was performed with adaptive focus control activated, 1024x1024 pixels, 200 Hz line scan speed and 4-line averaging. Images were processed using ImageJ 1.53c unless stated otherwise (6,7).

### 6. Clathrin dynamics: time-lapse confocal imaging

EYFP-CLTA cells transfected in a 6-well glass-bottom plate were gently washed 1x with 1 mL prewarmed DPBS (Thermo, 14200067), placed in prewarmed $CO_2$-independent medium consisting of DMEM-HEPES (Gibco, 21063029) supplemented with 10% FBS (Sigma, F7524) and 0.1% Gentamycin (Carl Roth, 2475.1) and imaged under incubation at 37 °C and 5% $CO_2$. Continuous time-lapse imaging was performed for 20 minutes with an image acquisition every minute. Images were taken according to previously described acquisition settings in three z-planes starting at the ventral membrane with 2 consecutive slices each 0.3 µm higher. In this article, only the z-slice corresponding to the ventral membrane is shown, which is the lowest z-position where fluorescent signal reaches a maximum brightness for the clathrin structures of interest. Discontinuous time-lapse imaging was performed over a time span of 8 h according to previously described acquisition settings. The position of the cell was stored in the LAS X Navigator which was used to retrieve the same cell at consecutive timepoints. Between timepoints, the sample was transferred back to the humidified incubator at 37 °C and 5% $CO_2$. Temporal color-coded images (as shown in Fig 2 and Fig S2) were created using ImageJ 1.53g (6,7). First, drift was corrected based on the fluorescent channel using the 3D drift correction plugin with standard settings and detection of slow drifts, sub pixel drift correction and edge enhanced images enabled. Afterwards, a temporal color code was applied for the fluorescent channel using the Temporal-Color code command that was adapted as suggested on the ImageJ forum (temporal color code bug 41323).

7. Super-resolution PALM-TIRF fluorescence microscopy

mEos3.2-CLTA samples transfected in a 35 mm disk were gently washed 1x with 1 mL prewarmed DPBS and placed in 1 mL prewarmed HBSS (Gibco, 14025100). For retrieval of the same cell in confocal and PALM-TIRF microscopy, mEos3.2-CLTA samples in a 35 mm disk with imprinted grid were first imaged in confocal microscopy as previously described, the same cell was retrieved using the grid coordinates and imaged on the PALM-TIRF microscope immediately afterwards. PALM-TIRF imaging was performed on a home-build fluorescence widefield microscope with built-in through-the-objective total internal reflection (TIRF) illumination. This IX83 inverted microscope (Olympus IX83 frame S1F-3, Olympus Optical, Tokyo, Japan) has been previously described by Rocha et al. (9) and includes dichroic mirrors to combine the laser lines, neutral density filters to adjust the laser intensity (Newport Corporation, Irvine, CA, USA), a 10x beam expander (Linos, Qioptiq, Luxembourg), a focusing lens to enable TIRF illumination (f¼ 500 mm, BK 7, Newport Corporation), a 60x TIRF oil objective (Olympus Optical, NA: 1.45), a z488/561 dichroic mirror to separate excitation and emission, and a 2.5x projection lens (Olympus) placed before a 512x512 pixels EM-CCD camera (ImagEM, Hamamatsu Photonics, Hamamatsu, Japan) with a physical pixel size of approximately 107 nm.

For image acquisition, first a diffraction-limited image was taken by 488 nm excitation (200 mW 488 nm diode laser, Sapphire, Coherent) that passed a series of neutral density filters (1.62 OD) before reaching the sample, while emission was captured between 500-550 nm (HQ525/50 band-pass filter, Chroma Technology). For PALM imaging, mEos3.2 was stochastically activated using a 405 nm diode switching laser (100 mW laser Cube Coherent Inc., Santa Clara, CA, USA used at 3-6% intensity with neutral density filters 2.12 OD, resulting in an effective laser power of 0.46-2.70 µW at the objective). Stochastically activated fluorophores were excited with a 561 nm laser (200 mW laser, Sapphire, Coherent Inc. with neutral density filters 0.08 OD) and emission was captured between 575-615 nm (HQ595/40 band pass filter, Chroma Technology). Image acquisition was performed by recording 2 x 5000 frames with an exposure time of 0.030530 s/frame. Transmission images were taken before, in between and after the 2 x 5000 frames to ensure the sample did not suffer from laser illumination during the imaging procedure.

For image reconstruction, a custom-written MATLAB algorithm (MATLAB R2022a) based on the open-source Localizer software package (10) was employed to localize the center positions of the emitters in each of the 10 000 frames by fitting them with a 2D Gaussian function with PSF standard deviation factor 1.8 and intensity selection sigma factor 25. By merging these retrieved positions, a super-resolved image with a lateral resolution of approximately 30-40 nm was obtained.

8. Clathrin topology quantification: cluster identification and classification in PALM-TIRF images

Clathrin topology was quantified in a PALM-TIRF dataset using a two-step MATLAB analysis that involved identification and classification of individual clathrin clusters. For identification of individual clathrin clusters, the DBSCAN data clustering algorithm was employed. First, images were gated to exclude background localizations falling outside the boundaries of the cell, which was defined by grouping nearby localizations into a region and selecting the largest region in the field of view. Then, the MATLAB DBSCAN function (11) was executed for a minimum of 17 neighbors (MinPts) within a search radius (ε) of 0.55. These parameters were optimized based on visual observations on 6 different cellular regions from both poorly and highly metastatic cells by testing all possible combinations for MinPts = 8 – 23 and ε = 0.35 – 0.65 (Fig S6A for example). Since this approach identified also many extremely small clusters lying predominantly in noisy regions that correspond to single clathrin molecules rather than a CCS, only clusters with ≥ 50 localizations were retained for further analysis (Fig S6B).

To distinguish classical from alternative CCSs, identified clusters with ≥ 50 localizations were classified using a custom-written MATLAB algorithm. This identification was based on characteristics of individual clusters, more specifically the cluster area, the number of localizations within the cluster (NrPts, associated with cluster brightness), the cluster eccentricity (associated with cluster roundness), the cluster perimeter and the distance to the nearest neighboring cluster (MinDistance). Cluster eccentricity was calculated using the open-source MinVolEllipse function (12). The classification model stated that if:

$$\frac{Area * \left(\frac{NrPts}{1000}\right) * \ Eccentricity * \ Perimeter}{MinDistance} > P90\ (classical\ CCSs)$$

the cluster was considered as an FCL, otherwise the cluster was classified as a classical CCS. The location of each of these terms in the numerator or denominator was carefully considered. Namely, a non-classical FCL is, from the literature or from our own observations (Fig S6C), expected to have a larger surface area, be brighter (larger NrPts), have a more elongated shape (eccentricity closer to unity), have a more irregular shape (larger perimeter) and would lie closer to other clusters (smaller MinDistance). Therefore, the model would yield a higher numerical output for an alternative FCL than for a classical CCS. This output was compared to a global threshold value (P90 classical CCSs) that was deduced from 1247 classical CCSs and 1042 FCLs that we manually selected from 3-5 different cellular regions containing predominantly classical CCSs and FCLs, respectively. The global threshold was set so that maximally 10% of the classical CCSs were incorrectly classified. This threshold was employed for quantification of both the KM12 and the SW model. The model was validated by visual confirmation to correctly identify the different structures. Cells that were too dim or too bright to allow for proper cluster identification and classification were excluded from the dataset.

Cluster classification of the PALM-TIRF dataset for KM12 and SW cell models identified on average 429 CCSs per cell, adding to a total average of 15 003 CCSs per cell line. The percentage classical CCSs and alternative FCLs identified in every cell, organized per cell line, can be found in Fig S7.

9. Clathrin topology quantification: post-classification processing and statistics

The above-described model resulted in a list of identified CCSs for every cell, in which each cluster was classified either as a classical CCS or as an alternative FCL. The individual cluster properties that were employed for the cluster classification model were also listed for every cluster.

Firstly, for analysis of individual cluster area, the median area of all classical CCSs and the median area of all alternative FCLs were calculated per cell. Considering the variance between different measured cells, these medians were considered as datapoints in our analysis (Fig S6E). Medians were pooled per cell line while keeping classical CCS and FCL categories separated, and they were plotted and analyzed according to these categories. Secondly, to check whether cluster area depends on cell area (Fig S6F), individual cluster area was plotted in function of cell area for all the pooled measured cells while only distinguishing between classical CCS and alternative FCL categories. Thirdly, the ratio of (number FCLs per number classical CCSs) (Fig 4B) was calculated for each of the measured cells. Each ratio was considered as a datapoint in our analysis, they were pooled per cell line, plotted and analyzed according to these categories. Lastly, the percentage classical CCSs and percentage alternative FCLs (Fig S7) were calculated per cell, by taking the ratio of number classical CCSs or number of FCLs divided by the total number of identified clathrin-coated structures, respectively. Each ratio calculated per cell was considered as a datapoint in our analysis, they were pooled per cell line, plotted and analyzed according to these categories. Statistical analyses were performed using the open-source PlotsOfDifferences web app that determines statistical significances based on randomization of the data and therefore does not assume a specific distribution of the data (13). Statistical significances were evaluated for $p > 0.05$ (ns), $p \leq 0.05$ (*), $p \leq 0.01$ (**) and $p \leq 0.001$ (***).

### Supplementary Info References


1. Holger Hoffmann (2022). Violin Plot (https://www.mathworks.com/matlabcentral/fileexchange/45134-violin-plot), MATLAB Central File Exchange. Retrieved October 21, 2022.
2. Morikawa K, Walker SM, Jessup JM, Fidler I. In vivo selection of highly metastatic cells from surgical specimens of different primary human colon carcinomas implanted into nude mice. Cancer Res. 1988;48(7):1943–8.
3. Morikawa K, Walker SM, Nakajima M, Pathak S, Jessup JM, Fidler I. Influence of organ environment on the growth, selection, and metastasis of human colon carcinoma cells in nude mice. Cancer Res. 1988;48(23):6863–71.
4. Wiśniewski JR, Gaugaz FZ. Fast and sensitive total protein and peptide assays for proteomic analysis. Anal Chem. 2015;87(8):4110–6.
5. Solís-Fernández G, Montero-Calle A, Martínez-Useros J, López-Janeiro Á, Ríos VDL, Sanz R, et al. Spatial Proteomic Analysis of Isogenic Metastatic Colorectal Cancer Cells Reveals Key Dysregulated Proteins Associated with Lymph Node, Liver, and Lung Metastasis. Cells. 2022;11(447):1–23.
6. Abràmoff MD, Magalhães PJ, Ram SJ. Image processing with imageJ. Biophotonics Int. 2004;11(7):36–41.
7. Schneider CA, Rasband WS, Eliceiri KW. NIH Image to ImageJ: 25 years of image analysis. Nat Methods. 2012;9(7):671–5.
8. Rust MJ, Lakadamyali M, Zhang F, Zhuang X. Assembly of endocytic machinery around individual influenza viruses during viral entry. Nat Struct Mol Biol. 2004;11(6):567–73.
9. Rocha S, De Keersmaecker H, Uji-i H, Hofkens J, Mizuno H. Photoswitchable fluorescent proteins for superresolution fluorescence microscopy circumventing the diffraction limit of light. In: Engelborghs, Y, Visser, A (eds) Fluorescence Spectroscopy and Microscopy Methods in Molecular Biology, vol 1076 Humana Press, Totowa, NJ. 2014.
10. Dedecker P, Duwé S, Neely RK, Zhanbg J. Localizer: fast, accurate, open-source, and modular software package for superresolution microscopy. J Biomed Opt. 2012;17(12):126008-1–5.
11. Ester M, Kriegel H-P, Sander J, Xu X. A density-based algorithm for discovering clusters in large spatial databases with noise. KDD-96 Proc. 1996;226–31.
12. Nima Moshtagh (2022). Minimum Volume Enclosing Ellipsoid (https://www.mathworks.com/matlabcentral/fileexchange/9542-minimum-volume-enclosing-ellipsoid), MATLAB Central File Exchange. Retrieved October 21, 2022.
13. Goedhart J. PlotsOfDifferences - a web app for the quantitative comparison of unpaired data. BioRxiv Prepr. 2019;1–7.
